# Verbal Episodic Processing in Newborns

**DOI:** 10.1101/2025.09.19.677368

**Authors:** Emma Visibelli, Ana Fló, Eugenio Baraldi, Silvia Benavides-Varela

**Affiliations:** Padova Neuroscience Center | University of Padua, Italy, 35131; Department of Developmental Psychology and Socialization | University of Padua, Italy, 35131; Department of Women’s and Children’s Health, Neonatal Intensive Care Unit | University Hospital of Padova, Padova, Italy, 35128

**Keywords:** Verbal memory, acoustic variability, speaker identity, newborns, functional near-infrared spectroscopy

## Abstract

During the first period of life, human infants rapidly and effortlessly acquire the languages they are exposed to. Although memory is central to this process, the nature of early verbal memory systems and the factors that determine retention and forgetting remain largely unknown. Behavioural and brain measures have demonstrated memory formation in newborns. However, word traces fade in the face of acoustic overlap, leading to interference and forgetting. Here, we investigate whether speakers’ identity changes facilitate the separation into distinct acoustic episodes and the creation of non-overlapping verbal memories. Newborns (0-4 days-old) were tested in a familiarization-interference-test protocol, while neural cortical activity was recorded using functional Near-Infrared Spectroscopy (fNIRS). The results showed higher neural activation to novel words than to familiar ones during the test phase, indicating that the infants recognized the familiar words despite potentially interfering sounds. The recognition response was measured over the left inferior frontal gyrus (IFG) and superior temporal gyrus (STG) areas known to be crucial for encoding auditory information and language processing. The neural response also included the right IFG and STG, involved in interpreting vocal social cues and speaker recognition. The results indicate that speaker identity is a key feature in the formation of verbal memories from birth, facilitating separability, possibly through early source–content binding (i.e., what–who), a precursor to fully mature episodic memory.

**Impact Statement:** Speaker identity is a distinguishing feature at birth and highlights the episodic nature of humans’ first-stored verbal memories.

## Introduction

Word recognition entails processing and integrating various linguistic features, such as phonological content, along with contextual or indexical information, like speaker identity, accent, and emotional content, which are crucial for communication. Theoretical approaches to speech representation hold contrasting views on the role of indexical features in word recognition. Abstractionist models assumed that variability needs to be normalized or stripped away so that speech sounds could be recognized (e.g., Halle, 1985; McClelland & Elman, 1986; Norris et al., 2000; Pisoni & Luce, 1987). Episodic or exemplar approaches adopt an alternative perspective, assuming that memories of linguistic utterances are bound to indexical information (e.g., (Goldinger, 1996; Nygaard et al., 1994; Palmeri et al., 1993). The balance between forming exemplar memories and creating normalized word prototypes is crucial during language acquisition. Indexical information may aid in distinguishing memories, while abstract representations are necessary for generalization. However, how infants encode language as they develop is still not well understood.

When encoding word forms, young infants remember not just the words themselves but also specific indexical properties such as the speaker (Houston & Jusczyk, 2000), stress, amplitude, and affect (Singh et al., 2004; see Van Heugten et al., 2015). However, their learning is context-dependent: low-variability conditions promote the learning of specific examples (Houston & Jusczyk, 2000; Jusczyk & Aslin, 1995; Singh et al., 2004) and high-variability conditions facilitate the learning of abstract word prototypes (Singh, 2008). Current models of infant language comprehension (Jusczyk, 1997; Werker & Curtin, 2005) propose that in early stages, infants match specific sounds to stored instances of words and subsequently generate abstract word prototypes. In line with this, we hypothesize that speaker changes play a critical role in verbal memories’ formation at birth by providing indexical information for memory separation.

Verbal memory formation at birth is not well understood. Vast research on language processing supports the storage of both linguistic and speaker-specific information in newborns. Neonates readily distinguish phonetic changes (Cheour-Luhtanen et al., 1995; Dehaene-Lambertz & Pena, 2001), extract words from continuous speech (Fló et al., 2019, 2022), and detect speech structure (Benavides-Varela & Gervain, 2017; Gervain et al., 2008; Martinez-Alvarez et al., 2023), even amidst variability in speakers (Fló et al., 2025; Mahmoudzadeh et al., 2013a). Newborns also react to indexical features such as between-accent differences (Giordano et al., 2021) and are particularly sensitive to familiar voices (DeCasper & Fifer, 1980; Mehler et al., 1978; Spence & Freeman, 1996). Moreover, phonological processing is lateralized to the left hemisphere, while voice-related information shows right lateralization already in young infants (Blasi et al., 2011; Spence & Freeman, 1996; see review Grossmann et al., 2010). While these findings support normalized phonological representations and parallel processing of phonological and contextual features, it remains unclear how these features are integrated to form verbal memories at birth and how they can determine memory formation or forgetting.

Benavides-Varela and colleagues used functional near-infrared spectroscopy (fNIRS) to investigate the formation of word memories at birth, including the brain areas supporting this cognitive capacity and the factors that determine their loss or retention. The authors found that newborns familiarized with a 2-syllable word sound (hereafter referred to as word) show a recognition response after a few-minute-long retention period, which was characterized by decreased activity towards the familiar word and increased response to a novel word over temporal, frontal, and parietal areas (Benavides-Varela, 2012; Benavides-Varela et al., 2011, 2012). This research also indicated that under some circumstances, newborns’ memories appear fragile and highly vulnerable to interference. For example, recognition does not persist when neonates hear another word produced by the same speaker during the retention period. Interestingly, unlike speech, instrumental music presented during this retention phase does not interfere with the familiar memory trace (Benavides-Varela et al., 2011). The phenomenon could be partly explained by retroactive interference, which occurs when novel information disrupts the retention of previously learned items (Müller & Pilzecker, 1900). One factor that may influence retroactive interference is the degree of neural overlap between the information to be encoded and the interfering stimuli. Since instrumental music and speech processing recruit partially distinct brain areas (in adults: Peretz et al., 2015; Zatorre et al., 2002; infants: Dehaene-Lambertz et al., 2002; and newborns: Kotilahti et al., 2010; Perani et al., 2010), this could explain the absence of music-speech interference. However, if this were the sole factor determining interference, speech-speech retroactive interference would render language learning impossible in real-life conditions. Here, we propose a complementary explanation for the retroactive interference described in previous studies: various features may be integrated to assess the similarities or differences between two auditory events, facilitating the separability of newly arriving information and, therefore, memory storage. Specifically, non-phonological information in speech, such as a speaker change, could serve as indexical information—acting as markers that signify the end of one event and the beginning of another—thereby facilitating the contrast and separability of verbal memories early in life. According to this hypothesis, the presence of speech during the retention period will not always lead to forgetting.

To test our hypothesis, we implemented a protocol derived from the work of Benavides-Varela et al. (2011, 2012). Newborns were first familiarized with a pseudoword produced by a single speaker. Immediately after, they were exposed to an interfering word. Then, in the test, the familiarization word or a completely novel word was presented. Like in Benavides-Varela et al., the interfering, the familiar, and the novel words had similar intensity, duration, pitch, syllable structure, etc. (see Supplementary Table 1 in Supplementary Materials (SI)). Instead, the familiarization was reduced from ten to five blocks, and the retention interval increased from two to three minutes, making the paradigm more challenging. These methodological adjustments allow for a meaningful comparison with previous studies: if newborns forgot the familiarization word when an interfering word was presented as in Benavides-Varela et al. (2011), they are expected to forget it also under our more challenging paradigm. However, unlike the previous work, the interfering word here was uttered by a different speaker. We hypothesize that if the voice distinction promotes memory separation, there should be a differential hemodynamic response between the familiar and a novel word in the test phase, signalling recognition. Instead, a failure in word recognition would reveal that a voice change is not sufficient to overcome the interference effect previously reported with this paradigm. We included 32 neonates in the final analysis, a number comparable to the 28 infants tested in the previous experiment showing interference (Benavides-Varela et al., 2011), which should warrant enough statistical power. A remarkable difference between this and previous studies is the use of a within-subject design with two familiarization-interference-test sequences (one testing responses to novel words and the other to familiar words). This design controls for differences in anatomy, physiology, and brain activity across individuals while increasing statistical power.

## Results

In this paradigm, responses are expected to change over time due to habituation and recognition dynamics. Accordingly, it is not appropriate to average responses across blocks belonging to the familiarization and test phases. Block-level analyses were thus conducted using Linear Mixed Models (LMM), which are well-suited to handle missing values. This approach was necessary because each subject provides a unique instance for each block, which inevitably leads to missing values in the dataset— for example, when a motion artifact renders an entire block invalid for that subject. We used the hemodynamic response over each block and the six Regions of Interest (ROIs) covered by the probe as the dependent variable. We decided to analyse the data at the ROI level, since channel-level analysis is potentially more susceptible to optodes placement differences. Moreover, channel-level analysis increases the number of comparisons in a protocol that already needs to compare activation over multiple blocks. Nevertheless, analysing the data at the channel level yielded similar results (see Supplementary Table 2). The ROIs were symmetric between hemispheres and included the inferior frontal gyrus left and right (IFGl, IFGr), the superior temporal gyrus left and right (STGl, STGr), and the parietal lobe left and right (PLl, PLr) (Figure 1A). We modelled fixed effects (e.g., condition: *same* or *novel*) nested within the block number and the ROIs, while including participants as random effects. Each such model indicates whether there are significant fixed effects within each block and ROI, without the need to correct for multiple comparisons for the number of blocks and ROIs. Only results for oxy-haemoglobin (HbO) are presented here. Results for deoxy-haemoglobin (HbR) were less clear and are presented in the SI (Supplementary Figure 2).

**Figure 1.**
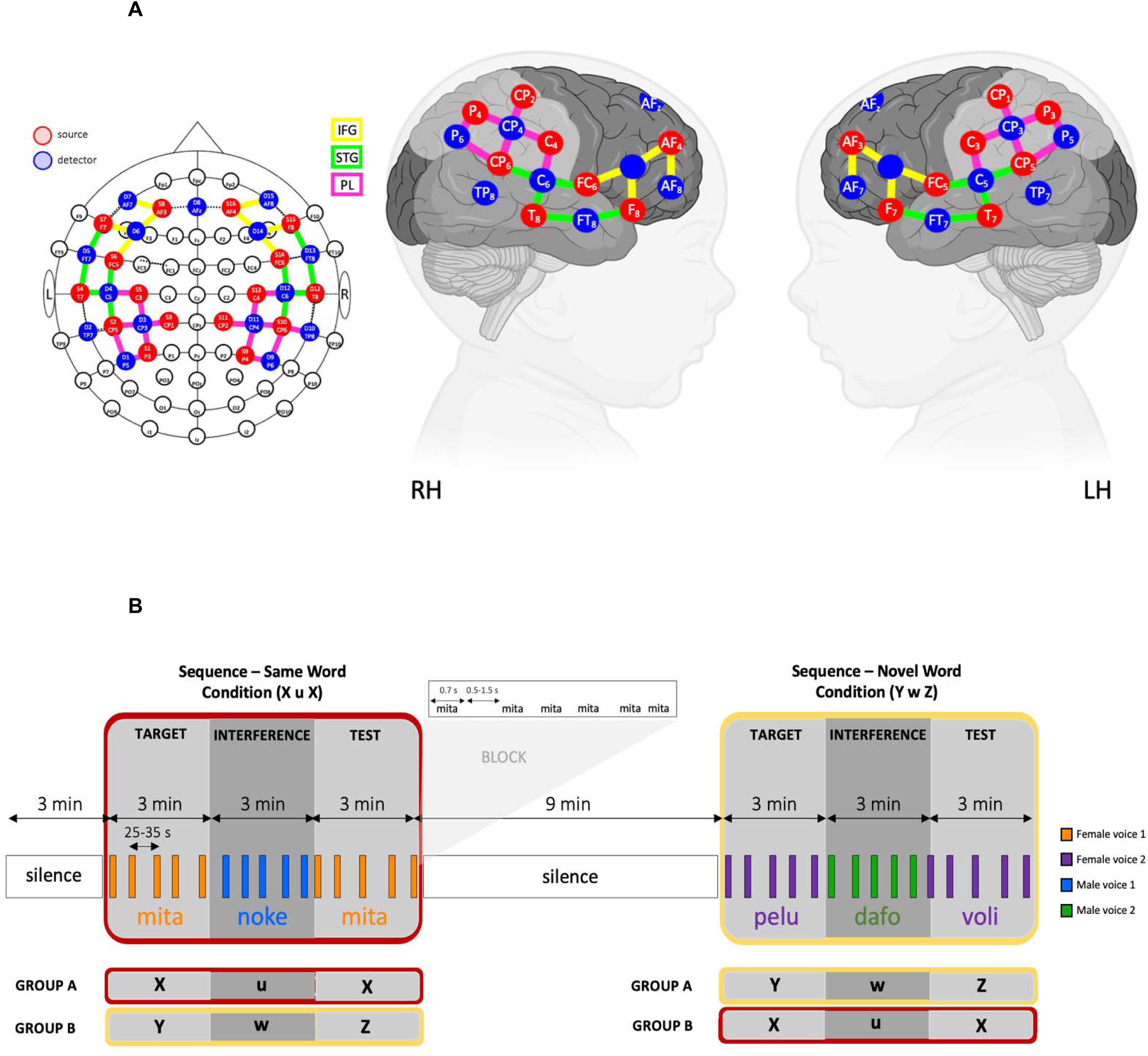
Experimental protocol. **(A)** Illustrative 42-channel fNIRS Montage. S (red) = source, D (blue) = detector. Placement indicated using the 10-10 standard EEG system. Regions of interest are indicated in yellow = inferior frontal gyrus (IFG), in green = superior temporal gyrus (STG), and in pink = parietal lobes (PL). **(B)** Familiarization-interference-test paradigm. Each subject was tested in two sequences separated by 9 minutes of silence: in one sequence, newborns heard the same word during familiarization and test (same-word condition; X u X), and in the other sequence, a novel word was presented during the test phase (novel-word condition; Y w Z). The order of the conditions, the words and the voices used in the different phases were counterbalanced across participants.

### Activity during familiarization

To assess potential habituation and novelty effects commonly observed in fNIRS data, we first tested whether the activity differed from zero by fitting the LMM *act ∼-1+block:ROI+(1|sub)* during the familiarization blocks. This model provides one coefficient for each ROI and block (β*(ROI_j_, block_i_)*) representing the activation. The model showed a positive activation in block 2 within left IFG (β*(IFGl, b2)=0.194, SE=0.064, p = 0.024*) and during blocks 4 and 5 within left STG (β*(STGl, b4)=0.173, SE=0.065, p = 0.008;* β*(STGl, b5)=0.128, SE=0.063, p = 0.044*) (Figure 2A).

**Figure 2.**
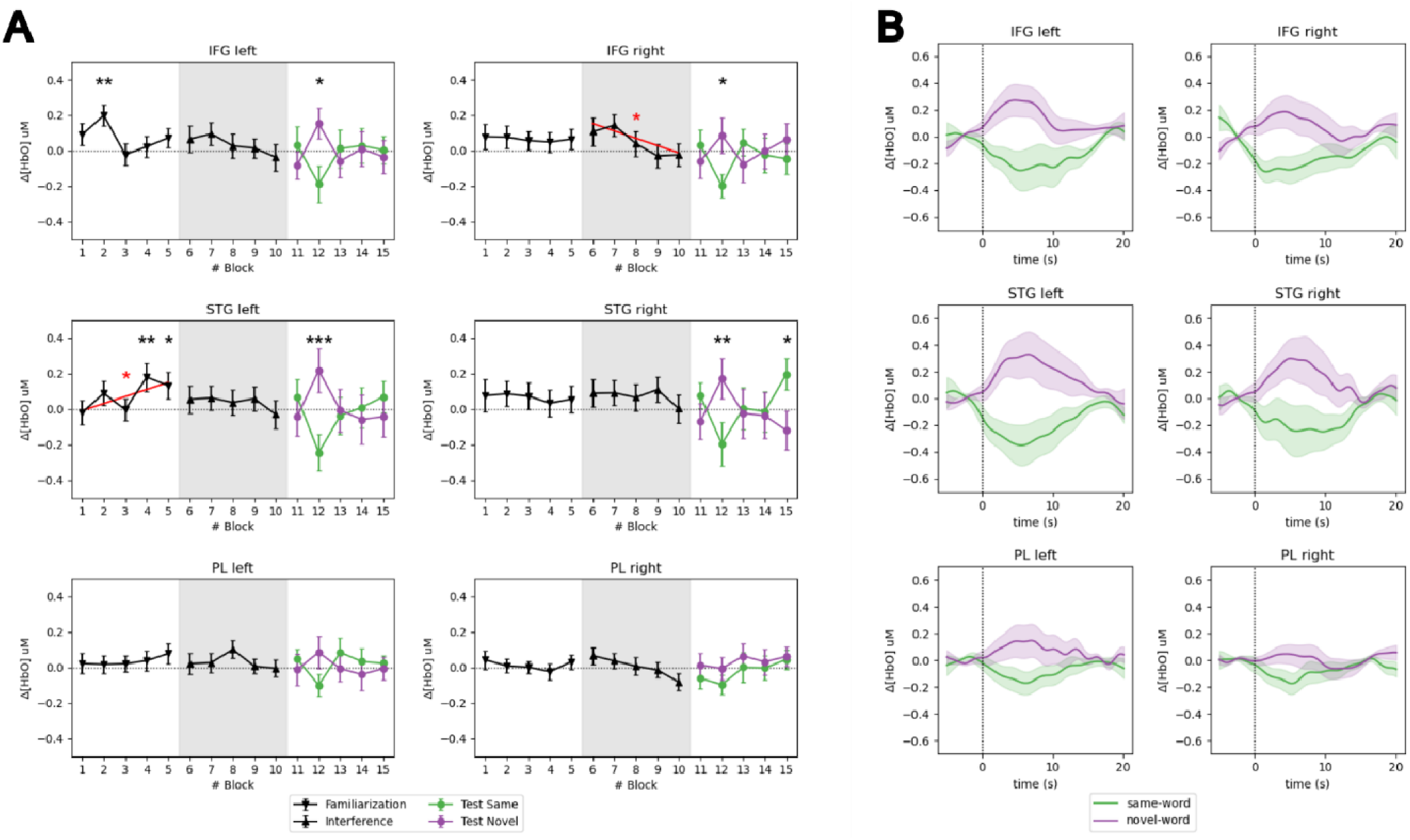
Standard recognition response with decreased activity for the familiar words and increased activity for the novel words in the test phase. **(A)** Mean activity for HbO per block during the familiarization, interference, and test phases. Error bars represent the standard errors. The black continuous line depicts responses averaged across all participants and conditions. The same-word condition (green) and the novel-word condition (purple) are plotted in the test phase. The black asterisks during the familiarization and interference phases indicate that the response differed from zero. The red lines indicate a significant linear trend, as indicated by the red asterisks. Black asterisks during the test phase indicate significant differences between conditions. **(B)** HRFs for HbO during the second block of the test phase, when relevant differences were observed between conditions. Shaded areas represent the standard error.

Additionally, we tested for linear changes in activity by fitting the LMM: *act ∼-1+ROI+ROI:blocknumber+(1|sub)*, with *blocknumber* coded from 0 to 4. This coding scheme allows the intercept term for each ROI to represent activity in the first block, while the corresponding slope term captures any linear change in activity across blocks.

The model showed a significant intercept in the left IFG (intercept=0.1105, SE=0.0535, p=0.040), indicating an initial positive activation and a significant positive slope in the left STG (slope=0.0396, SE=0.018, p=0.029), denoting a sustained increase in activity in this area (Figure 2A). An analogue analysis for the interference phase is presented in the SI (Supplementary Figure 4).

### Word recognition

We assessed recognition responses in the test phase by testing whether the activation pattern differed between the familiar and novel words. We employed an LMM, including condition as a fixed factor nested within the ROIs and blocks of the test phase *act∼-1+block:ROI+block:ROI:condition+(1|sub)*. Such a model provides, for each ROI and block, one coefficient quantifying activation in one condition and another quantifying the difference between conditions – thus, crucial for evaluating word recognition. The model showed a significantly higher activation during the second block of the test phase for the *novel-word* than the *same-word* condition over IFG and STG (β*(IFGl, b2)=0.322, SE=0.133, p = 0.015;* β*(IFGr, b2)=0.265, SE=0.133, p = 0.045;* β*(STGl, b2)=0.443, SE=0.133, p = 0.0009;* β*(STGr, b2)=0.348, SE=0.133, p = 0.009*). Instead, activity was higher for the *same-word* than the *novel-word* condition in the fifth block over STG right (β*(STGr, b5)=-0.320, SE=0.127, p = 0.012*). To investigate the presence of hemispheric differences in the main effect of condition revealed by the primary analysis, we ran an LMM restricted to the second block and the IFG and the STG separately (*act ∼cond*hemisphere+ (1 | sub)*). We found no significant effects of hemisphere or interaction, neither over IFG nor on STG (p>0.1) (see Figure 2A-B).

### Effects of the sequences order

In our within-subject design, group A first completed the same-word condition (X u X) and later the novel-word condition (Y w Z), while group B did the opposite (Figure 1B). Thus, the first sequence might influence the processing of the second sequence, potentially leading to differences between sequences and groups.

We looked for differences during the familiarization and interference phases by fitting an LMM contrasting (1) first and second sequence, (2) groups within the first sequence, and (3) groups within the second sequence. The contrasts were nested within blocks and ROIs, such that, for each ROI and block, a coefficient was fitted for each contrast (see details in SI). The model showed higher activation during the second than the first familiarization in the first block over IFG and STG (β*(IFGl, b1, contrast 1)=-0.254, SE=0.191, p = 0.033;* β*(STGl, b1, contrast 1)=-0.249, SE=0.191, p = 0.036;* β*(STGr, b1, contrast 1)=-0.322, SE=0.191, p = 0.0069*) (Supplementary Figure 3). Differences between groups were weak and restricted to higher activation in group B than A in the first block of the first sequence over the left STG (β*(STGl, b1, contrast 2)=-0.368, SE=0.182, p = 0.043*) and on the first block of the second sequence over the right STG (β*(STGr, b2, contrast 3)=-0.355, SE=0.175, p = 0.042*). Considering the small number of data points per group and sequence, these differences are likely due to noise. See in SI the analysis for the interference phase (Supplementary Figure 4).

Given the differences in activation between the first and second familiarization phases, we quantify linear changes in activity as we did previously, but separately for each familiarization sequence. For the first familiarization, the model showed a significant increase in activity in the left and right STG and left PL (p<0.05), while during the second familiarization, the activity was higher than zero in the first block and decreased with block number on the right STG and IFG (p<0.05) (detailed results are presented in Supplementary Figure 3).

To check for differences between the two groups during the testing phase, we fitted an LMM contrasting (1) the *same-word* and *novel-word* conditions, (2) the groups within the *same-word* condition (i.e., *same-word* presented in group A, thus, sequence 1, or group B, thus, sequence 2), and (3) the groups within the *novel-word* condition (i.e., *novel-word* presented in group A, sequence 2, or group B, thus, sequence 1). The contrasts were nested within blocks and ROIs, yielding a coefficient for each contrast within each ROI and block. In agreement with the overall results obtained when merging the two groups, the model showed a significant main effect of condition during the second block over IFG and STG (β*(IFGl, b2, contrast 1)=0.334, SE=0.131, p = 0.011;* β*(IFGr, b2, contrast 1)=0.293, SE=0.131, p = 0.026;* β*(STGl, b2, contrast 1)=0.459, SE=0.131, p = 0.00050;* β*(STGr, b2, contrast 1)=0.367, SE=0.131, p = 0.0053*), and during the fifth block over right STG (β*(STGr, b5, contrast 1)=-0.358, SE=0.128, p = 0.0051*). No significant differences were observed between groups (sequences) for the *same-word* condition (p> 0.05). However, the model showed significant group differences for the *novel-word*. Activation was higher for the *novel-word* in group A (*novel-word* in sequence 2) than in group B (*novel-word* in sequence 1) in the second block over right IFG (β*(IFGr, b2, contrast 3)=0.538, SE=0.183, p = 0.0031*) and in the third block over left and right STG (β*(STGl, b3, contrast 3)=0.394, SE=0.182, p = 0.031; STGr:* β*=0.533, SE=0.182, p = 0.0036*). Instead, activity for the *novel-word* was higher for group B than A in the fourth block over IFG and STG (β*(IFGl, b4, contrast 3)=-0.443, SE=0.198, p = 0.025;* β*(IFGr, b4, contrast 3)=-0.390, SE=0.198, p = 0.049;* β*(STGr, b4, contrast 3)=-0.403, SE=0.198, p = 0.042*).

This analysis confirms that both groups show a consistent recognition response (higher activity for the *novel-word* than the *same-word*) in the second block, over left and right STG, and left IFG. In addition, it indicates a more complex pattern of activation in the right IFG, with an interaction between condition and group. To better understand the effect, we performed Tukey’s multiple-comparison test. Results showed higher activation for *novel-word* in group A than all the other conditions (*novel-word* in group B: p=0.007, *same-word* in group A: p=0.0096, and *same-word* in group B: p=0.0073), confirming that the difference between conditions on the right IFG during the second block was driven by group A (Figure 3).

**Figure 3.**
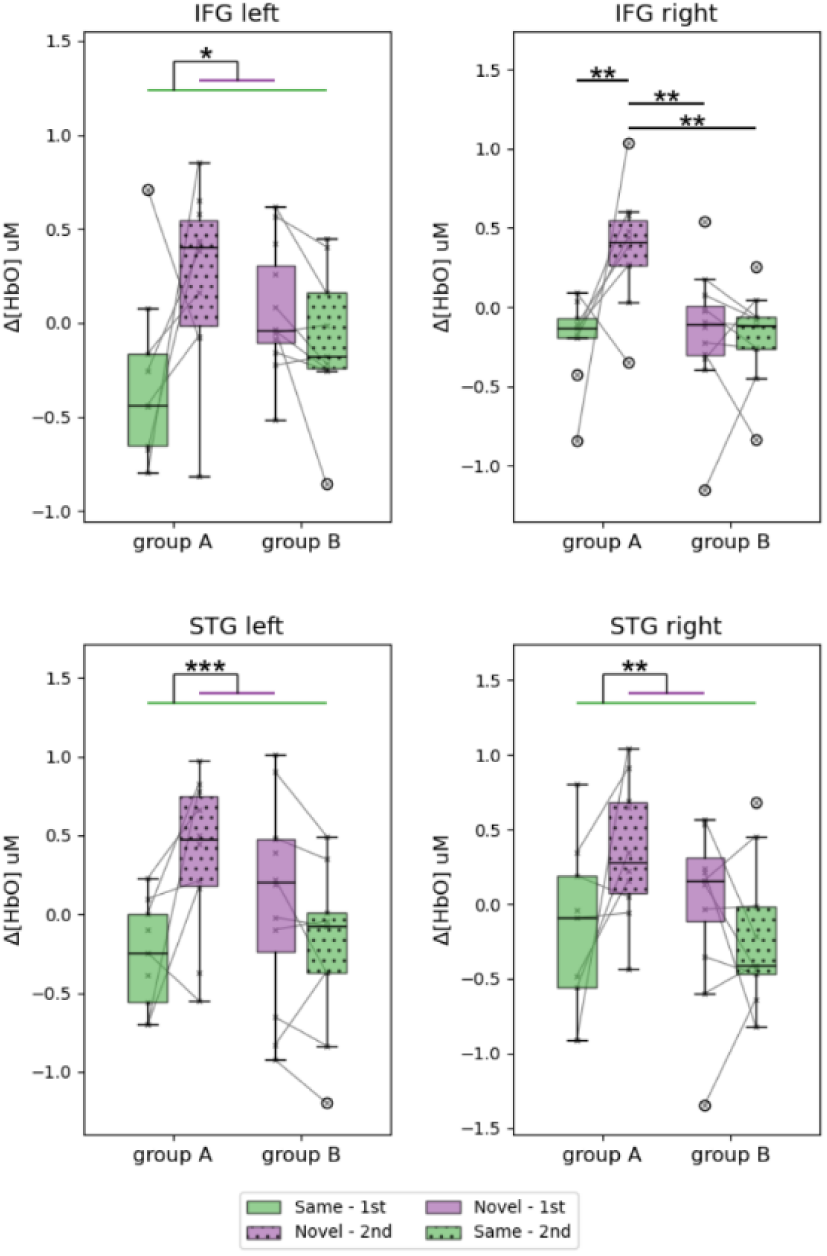
Differences in the response across groups during the second test block. Boxplots represent the mean HbO activity during the second block of the test phase, separated by condition (same=green, novel=purple), group (A or B), and sequence (first=full pattern or second=dotted pattern). Whiskers of the boxplot are defined based on 1.5 times the interquartile range, and data points outside these limits are plotted as circles. Asterisks indicated significant differences between conditions or groups. A significant effect of conditions was observed in the left IFG and the left and right STG, reflecting recognition in both groups. Instead, an interaction effect was present in the right IFG with higher activity in the *novel-word* condition for group A.

## Discussion

### The role of variability in early memory processes

In the current study, we investigated the conditions that promote the formation of separate memory traces of linguistic stimuli at birth. We observed a persistent neural signature of recognition, namely a differential response between the familiar and novel words, by introducing a change in the speaker uttering the interference word. In Benavides-Varela et al. (2011), word recognition in neonates vanished when they heard an interference word pronounced by the same speaker who uttered the familiarization word. We hypothesize that the shared voice feature could have increased perceived acoustic overlap (Apfelbaum & McMurray, 2011), thereby causing interference. In our study, the presence of a new speaker might rather act as a conspicuous cue signalling the beginning of a new acoustic episode and facilitating the separation of linguistic memory traces.

These results demonstrate that, under certain conditions, newborns can retain verbal memories even when the language networks continuously receive new verbal information, as in real life. These results are in line with episodic models of early speech perception, assuming that infants initially store words in an instance-specific fashion comprising both phonological details and speaker identity (Jusczyk, 1997; Werker & Curtin, 2005). Furthermore, the findings extend these models by offering empirical evidence of episodic encoding in newborns: early word-form representations are, at least to some extent, linked to the acoustic realization of the word and, when it comes to early signal-to-word form mappings, the newborn brain attributes significant relevance to voices. The speaker’s identity may thus represent a critical distinguishing factor essential for early communication and memory. Forgács et al. (2022) recently showed that alternation between female and male voices, combined with partial variability in the syllable stream, elicited greater activation in the left fronto-temporal regions. This finding suggests that the facilitation of verbal human memory in newborns might also be related to the heightened neural activation associated with communicative attribution. In this view, infants may interpret such vocal alternations as indicative of a communicative exchange, thereby enhancing their ability to segregate and store the pseudo-words presented as stimuli.

These findings speak to the relevance of certain cues in the sequential processing of speech input, but do not inform us about the possibility that newborns can handle indexical variation (i.e., speaker changes or changes in intonation and emotional content) during the presentation of the word in the familiarization phase or recognize the familiar word irrespective of possible indexical variations in the test. There are some hints in the literature suggesting that this might be the case. Newborns robustly encode words presented in concomitance with other words, suggesting that word-memories can be formed in the face of input variability (Benavides-Varela et al., 2012). Moreover, newborns show recognition of pseudowords despite prosodic differences (Fló et al., 2019) and compute regularities over phonetic content, disregarding the voice content (Fló et al., 2025). Thus, it is possible that if a variety of diverse tokens are presented during learning, a robust and generalizable representation could emerge as early as birth. While this question lies beyond the scope of the present study, it could provide additional insights into early word recognition processes.

### Signature of word recognition and areas recruited for memory retrieval

In the current study, we observed the typical recognition response characterized by an increase in activity for the novel word and a decrease for the familiar one, consistent with previous studies using a similar paradigm. Although the fNIRS system and optodes positioning slightly differed from those of previous studies (e.g., Benavides-Varela and colleagues’ system covered more prefrontal areas, whilst our configuration only reached the IFG), the activation pattern in the temporal and frontal areas is generally consistent across studies. In the present study, the effect was bilateral over the IFG and STG, known to play a crucial role in language processing and in interpreting vocal social cues in the left and right hemisphere respectively. In particular, left frontal regions, including the IFG, are associated with processing, retrieving, and manipulating phonological information (e.g., Bunge, 2001; Hickok & Poeppel, 2007; Novick et al., 2010; Thompson-Schill et al., 1997), and the left STG plays a crucial role in phonological and semantic processing (Hickok & Poeppel, 2007) by encoding fast temporal (phonetic) information (DeWitt & Rauschecker, 2012; Mesgarani et al., 2014; Zatorre & Belin, 2001) and integrating auditory information within verbal memory (Cabeza & Nyberg, 2000). Conversely, speaker recognition relies primarily on a right-lateralized network (Mathias & Von Kriegstein, 2014), with the right IFG and STG essential for processing prosody, rhythm, and vocal social cues such as emotional state and intent (Agus et al., 2017; Belin et al., 2000, 2002; Bodin et al., 2018; Fecteau et al., 2004; Pernet et al., 2015; Wildgruber et al., 2006; Zatorre et al., 2002).

Precursors of the same organization and hemispheric specialization seem to be in place early on in life (Dehaene-Lambertz & Baillet, 1998; Mahmoudzadeh et al., 2013b; Telkemeyer et al., 2009), including the activation of left fronto-temporal areas associated with language processing (Alexopoulos et al., 2021, 2022; Dehaene-Lambertz et al., 2002; Peña et al., 2003) and functional specialization of the right STS for voice processing (Blasi et al., 2011; Cheng et al., 2012; Grossmann et al., 2010; Schönwiesner et al., 2005; Simon et al., 2009). The different responses we observed between new and familiar words after a three-minute retention period align with the retrieval of the verbal memory. Therefore, the bilateral concurrent responses over the IFG and STG suggest that linguistic and non-linguistic features of the word contribute to the recognition response in this context.

### Timing of the response: word recognition in the second block of the test phase

Factors including experimental design and stimulus complexity are known to influence hemodynamic responses in newborns and infants across tasks (Issard & Gervain, 2018). In this paradigm, an interplay between familiarization length and the presence of interfering sounds might determine when the differential response between a novel and a familiar stimulus emerges. In simple experiments, recognition is detected in the first block of the test when a single identical word is repeated over 6 minutes in the familiarization and when no interfering sounds are presented during the retention interval (e.g., Benavides-Varela et al., 2011). In more complex designs, the recognition was delayed to the second block when an interfering word alternates with the to-be-remembered word during encoding (Benavides-Varela et al., 2012). Similarly, in the current study, the recognition response emerged in the second block of the test phase when an interfering word sound was presented during the retention phase. Thus, while newborns can recognize word sounds under complex conditions, facing these challenges influences the timing of the recognition response in the test phase, requiring additional cues or extended processing time for activation.

### Familiarization phase

Stable activity was registered with no obvious attenuation of the neural response over the three minutes in the familiarization phase. This general pattern was observed in most areas, but in the left STG, where the neural response showed repetition enhancement over time. Neural suppression (habituation) or enhancement, while expected in the context of repeated stimuli, is not consistently found across fNIRS studies in infants and newborns. Various factors may influence hemodynamic patterns over time. First, some studies using a protocol similar to ours found habituation over the left frontal areas in newborns when target words are presented in “ecological conditions”, that is, interleaved with other words (Benavides-Varela et al., 2012). By contrast, habituation is not reported when the familiarization is homogeneous (Benavides-Varela, 2012; Benavides-Varela et al., 2011), as in the current study. This suggests that the amount of information present during the learning phase modulates newborns’ fNIRS neural dynamics. The role of stimulus complexity has also been demonstrated in fNIRS studies of rule-learning in newborns. While highly variable speech sequences elicited left-lateralized repetition enhancement across blocks for ABB artificial grammar and no variations for ABC grammar (Gervain et al., 2008), simpler stimuli and presentation conditions (blocked rather than interleaved) evoked a stable response for the simpler ABB grammars and a repetition enhancement effect over time for ABC grammars (Bouchon et al., 2015). Second, methodological factors, such as the frequency and number of stimulus repetitions, are known to influence the habituation (Rankin et al., 2009). Thus, the sparse stimuli presentation typical of fNIRS block-designs (with stimuli followed by periods of 20-25s of silence), along with the reduced number of blocks employed in the present study, may have also contributed to the patterns observed. Third, Katus et al. (2023) recently tested habituation to a female voice in 1-month-old (asleep), 5-month-old (awake), and 18-month-old (awake) infants. They found that habituation began to emerge at five months and became strong by eighteen months. Similarly, another study revealed stronger effects of habituation in 8-month-old awake infants compared to 5-month-olds (Lloyd-Fox et al., 2019). Altogether, these studies show that developmental changes and sleep state influence habituation as measured by fNIRS. It is therefore likely that all these factors (i.e., stimulus variability, stimulus frequency, duration of familiarization, participants’ age, and behavioural state) modulated the responses observed in the current study. Future research should carefully control these variables to further explore their role in learning and memory formation at birth.

### Cross-phase associations

A question raised by the present findings concerns how neural activity during the familiarization, interference, and test phases may relate to one another. The current study was not designed to test specific hypotheses about cross-phase dependencies, and therefore any such integration must remain descriptive. Nevertheless, the patterns of activity varied across ROIs, suggesting that encoding- and retrieval-related processes may interact in a region-specific manner. One illustrative example is the left STG, which showed a marked increase in activity during familiarization and a differential response between familiar and novel words in the earlier test blocks, which is consistent with a relationship between responses across phases within this ROI. By contrast, other regions, while also exhibiting a differential response at test, did not show comparably pronounced or systematic changes during the familiarization phase. In this context, converging evidence from developmental fNIRS work illustrates how such cross-phase dependencies can be revealed using connectivity-based approaches. For example, Benavides-Varela et al. (2017) observed a habituation-like hemodynamic response during encoding in left-frontal regions accompanied by progressive interactions between temporal and left-frontal regions. These interactions then served the recognition response in right-frontal and right-parietal regions, with connectivity from temporal areas emerging selectively for familiar items. The present data, based on univariate activation patterns, do not allow us to establish a direct causal link between activity during familiarization and subsequent test responses. Future work could further characterize how information is distributed across regions during familiarization, interference, and test, and how these patterns contribute complementary information to subsequent retrieval. Such approaches may also help clarify whether and how activity in specific regions during encoding predicts later recognition responses.

A further issue that merits discussion concerns the absence of an increase in activity during the interference phase. At first glance, this might seem at odds with the presence of a robust differential response in the test phase. However, the test phase engages memory recognition processes that rely on comparison of the incoming stimuli with stored representations and may therefore elicit a more sustained and detectable hemodynamic response than a mere acoustic change. Accordingly, a plausible explanation for this pattern relates to the temporal dynamics of the hemodynamic response, which in newborns may be less sensitive to transient sensory novelty than to the functional demands of higher-level processing. This interpretation is consistent with developmental evidence showing clearer neural responses to novel speech streams at later ages than in younger infants (e.g., at 3 months in Nakano et al., 2009; at 5 and 18 months but not at 1 month in Katus et al., 2023), suggesting that the detectability of sensory novelty effects by means of fNIRS possibly increases with maturation.

### Habituation, recognition, and novelty detection differences between groups

When interpreting the patterns in the familiarization, it is important to consider baseline activity. This consideration is especially relevant in within-subject designs, as the responses in the second session might be influenced by what newborns experienced in the first session. Our analysis captured these effects by showing higher activity in the first block of the second familiarization sequence than in the first. These results likely reflect a novelty response since all participants in the second familiarization session heard a new speaker pronouncing a completely novel word. At the same time, provides evidence that newborns can retain information from the first session over a 9-minute silent pause, allowing them to compare previously experienced episodes with newly encountered ones. These baseline differences result in distinct patterns over time: initial stronger activity followed by attenuation over blocks in the second sequences, while significant enhancement of the hemodynamic response is observed in the first sequences (Supplementary Figure 3A).

The within-subjects design also offers a valuable opportunity to investigate the responses to familiar and novel words when infants first heard a familiar word at the test, followed by a novel one in the second sequence, or vice versa. Notably, the main novel/recognition response over left and right STG and left IFG was consistent across groups. In addition, a group modulation was observed over the right IFG: Group A (which encountered the novel word condition during the second testing sequence), showed a stronger response in the right IFG than Group B (which experienced it during the first testing sequence). This effect, although unexpected, could be explained by the number of phonological or speaker changes newborns experienced until the novel stimulus was presented. Indeed, while the novel word corresponds to the fourth change for newborns in group A, it constitutes the second one for participants in group B. Variability of the stimulus facilitates learning and induces significant increments in attentional arousal (Cooper & Aslin, 1989; Fernald & Kuhl, 1987; Trainor et al., 1997), which might be reflected in the greater reactivity to novel information observed in group A. While more data should be gathered to better understand this phenomenon, the localization of the differential response in the right-lateralized areas further indicates that it pertains to the processing of vocal cues.

### Limitations

Some methodological and theoretical considerations merit attention. First, the length of the paradigm may have introduced a sampling bias, as more vulnerable infants who could not tolerate longer recording periods may have been excluded, and it also increases the likelihood of state changes. Newborn physiology changes rapidly, with transitions occurring within minutes, and such fluctuations can occur even in shorter experimental paradigms. Given recent evidence highlighting the distinct roles of sleep states in long-term cognitive development and functional connectivity (Lee et al., 2020; Uchitel et al., 2023), accounting for behavioural and physiological states during functional recordings should be a priority for future research (see Bastianello et al., 2025 for a comparative approach of sleep measures in infants).

A second consideration is that the present design did not include a control condition in which the interference word was spoken by the same speaker. However, a previous study employing a similar paradigm found that recognition does not persist under these circumstances (Benavides-Varela et al., 2011). Since our protocol was more demanding, with shorter familiarization and longer retention, it is unlikely that a same-speaker interfering word would have yielded different results in our setting. Thus, due to practical challenges and ethical considerations associated with testing newborns, we did not include this condition. Incorporating such a condition in future studies could help further refine the interpretation of the interference effects observed in early memory formation.

## Conclusion

Understanding the mechanisms governing memory and the factors enhancing it is crucial for comprehending language development. This study assessed newborns’ ability to retain a combination of speech sounds in the presence of acoustically novel interference. The findings showed that acoustic variability promotes separate memory traces of linguistic content rather than fully interfering with them. The presence of a new speaker may thus signal a new acoustic episode and facilitate the separation of linguistic memory traces. This suggests that the ability to encode information about the speaker is a fundamental process, potentially rooted in early brain mechanisms of cognitive development. This observation carries relevant implications when considered in relation to theories of memory, and models of memory development (Alberini & Travaglia, 2017; Behm et al., 2025; Yates et al., 2025). Episodic memory is a multifaceted construct that, in its mature form, entails the ability to retrieve past events with contextual detail, typically involving autobiographical recollection and the integration of *what–where–when* information (Tulving, 1993). Our study does not aim to demonstrate the presence of a fully developed episodic memory system at birth, nor do we claim that newborns’ performance satisfies all hallmark criteria of mature episodic memory. We focused on sensitivity to speaker identity as a contextual dimension relevant to memory formation. Within this narrower sense, both, the patterns of activation and the localization of the response provide evidence for early source–content binding (i.e., *what–who*), which can be considered a foundational aspect of episodic-like processing. Following up on this foundational step, future studies may track the gradual integration of additional aspects (*i.e. where-when*), ultimately leading to the maturation of a fully functional human episodic memory system, and investigate the neural mechanisms underlying this process.

## Methods

### Participants

Healthy full-term human newborns from a normal pregnancy (i.e., with no pathologies, perinatal, or neurological complications attested) were tested. Selection criteria included gestational age (GA) 37-42 weeks (range [37+1, 41+1]), Apgar scores ≥ 8 in the fifth and tenth minutes, absence of cephalohematoma or other conditions that could possibly affect cortical hemodynamics, intact hearing, head diameter within 32.5–37.0 cm range, and weight ≥ 2.5 Kg. Neonates were recruited from Neonatal Care Unit of the Unit of Neonatology and the Obstetric Division of the University Hospital of Padova between May 2023 and September 2023. Informed consent for participation in the experiment was obtained from parents. The Ethics Committee for Clinical Research of the Province of Padova, Italy, approved the study. Thirty-two infants who provided good quality data were included in the study (18 females; age range [0, 4] days; mean weight 3.364 Kg, SD 0.308 Kg). Eleven additional neonates were tested but not included in the analyses due to fussiness (not even five blocks free of artefacts in at least one of the testing sequences) (n=4), bad quality signal (more than 15 channels out of 42 marked as non-functional) (n=6), and technical problems (n=1).

### Stimuli

Five pseudowords (CVCV structure, stressed on the first syllable) were used in the study (target and test words: /*mita*/, /*pelu*/, /*voli*/; interference words: /*noke*/, /*dafo*/). Two female speakers recorded the target/test words (/*mita*/, /*pelu*/, /*voli*/), while two male speakers recorded the interference words (/*noke*/, /*dafo*/). Pseudowords were edited using the open-source Praat software (Boersma & van Heuven, 2001) to have a mean intensity of 70 dB and a duration of 700 ms. Detailed acoustic information can be found in the SI (Supplementary Table 1).

### Procedure and data acquisition

Neonates were tested in a dimly lit hospital room while lying in their cribs (N=23) or mothers’ arms (N=9), in quiet rest or sleeping, to ensure their comfort and maintain an ecologically valid environment. Pseudowords were presented through two loudspeakers using the Psychopy software (Peirce et al., 2019), while fNIRS data were recorded using the NIRx NIRSPort system (light sources of 760 and 850 nm, maximum intensity 25 mW per fiber per wavelength). We designed a probe configuration with 16 sources and 15 detectors forming 42 channels. The optodes were positioned according to the 10-20 system, with locations selected using the devfOLD toolbox (Fu & Richards, 2021) to cover the IFG, STG, and PL (Figure 1A). The average distance between sources and detectors was 2.13 cm (range = [1.75, 2.62] cm, SD = 0.21 cm), and the sampling rate was 7.63 Hz.

The experiment consisted of a Familiarization phase, an Interference/Retention phase, and a Test phase. Each phase lasted 3 minutes and comprised five blocks. In each of the five blocks, six pseudowords were presented (inter-stimulus interval = 0.5-1.5 s; inter-block interval = 25-35 s) (Figure 1B). The same pseudoword was presented in each phase.

A within-subject design was implemented by having two testing sequences separated by 9 minutes of silence: in Sequence 1, neonates heard the same word during familiarization and test (same-word condition; X u X), while in Sequence 2, a novel word was presented during the test phase (novel-word condition; Y w Z). The speakers and pseudowords were completely different in the two sequences. The pseudowords used in the different phases and the speakers were counterbalanced across participants. The order of the sequences was also counterbalanced across participants, resulting in Group A, presented with Sequence 1 and then Sequence 2, and Group B, presented with Sequence 2, followed by Sequence 1. Participants were initially assigned in a counterbalanced way to Group A or Group B, balancing the number of participants who completed the experiment across groups. Due to signal quality and attrition, the final sample consisted of 17 infants in Group A and 15 infants in Group B.

The paradigm was a modified version of a previously used experimental protocol (Benavides-Varela, 2012; Benavides-Varela et al., 2011, 2012). The Familiarization phase was reduced from ten to five blocks based on previous data showing that five blocks already result in habituation (Benavides-Varela et al., 2012) and to accommodate the two sequences within a single testing session. In addition, the retention period was extended from two to three minutes.

## Data Processing and Analysis

### Preprocessing

The first steps of data pre-processing were performed using custom functions and functions of the Homer3 fNIRS package (https://openfnirs.org/software/homer/homer3/; (Huppert et al., 2009) in Matlab 2024a. We first converted intensity to optical density using the Homer3 function hmrR_Intensity2OD and detected motion artifacts on optical density using a custom function. In brief, a copy of the data was created, and band-pass filtered between 0.01 and 0.7 Hz. Then, the maximum change in sliding time windows of 2 s (time step one sample) was computed, and a relative rejection threshold was obtained for each channel as *thresh* = *q*_3_ + 2 x (*q*_3 –_ *q*_1_), where *q*_3_ is the third quartile of the maximum changes distribution and *q*_1_ the first quartile. Using relative thresholds results in a better trade-off between data recovery and artifact detection without needing to optimize the thresholds for each experiment and subject (Fló et al., 2019, 2022). Time windows with a maximum change above the threshold were rejected, obtaining a rejection/inclusion matrix (tIncCh_MotArt) of the same size as the data. The procedure was repeated thrice or until less than 0.5 % of the data was rejected. Finally, a mask of 1 s was applied to the rejected data.

We used three metrics for channel pruning (i.e., defining non-functional channels): signal saturation, signal-to-noise ratio (SNR), and Scalp Coupling Index (SCI); for each of them, a matrix of the size of the data containing the metric per channel and sample was obtained. The saturation matrix was computed, marking saturated samples per channel when the intensity was outside the range [10^-6^, 2.5]. The SNR was computed in sliding time windows (length 5 s, step 2.5 s) as 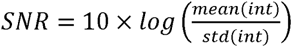, where *int* is the measured intensity. The matrix with the SNR was obtained based on the SNR in each time window. The SCI (Pollonini et al., 2014) was computed in sliding time windows (length 5 s, step 2.5 s) on the optical density band-pass filtered around the heartbeat frequency (heartbeat rate ± 0.4 Hz). The SCI matrix was then obtained. The heartbeat was estimated using the fNIRS recording in sliding time windows (length 60 s, step 15 s) as follows: the optical density was band pass filtered between 0.8 and 3.3 Hz, PCA was applied, and the autocorrelation was computed for the first principal component. Then, the cardiac frequency was estimated as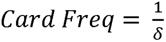, where *δ* is the time of the first peak of the autocorrelation –after the peak at zero-lag peak. The three metrics were evaluated on data segments free of motion artifacts for pruning channels (we call them *tInc_prunning*). *tInc_prunning* segments were defined as those with less than 30 % of the channels affected by motion artifacts and lasting at least 15 s (rejected segments shorter than 2 s were re-included). A channel was pruned if more than 30 % of the samples included in *tInc_prunning* showed: (1) saturation, (2) SNR<15, or (3) SCI<0.6. Subjects with more than 15 out of 42 channels pruned were excluded from the analysis.

Artifact correction techniques can reduce artifacts’ size, but no meaningful data can be recovered if the duration of the artifact is longer than an HRF. Since infants’ data might be contaminated with strong and long motion artifacts, we used the rejection matrix obtained from the artifacts detection step (*tIncCh_MotArt*) to define long segments heavily contaminated by motion and later reject blocks overlapping with them. These contaminated long-segments were defined as samples with more than 50% of channels contaminated with motion artifacts and lasting at least 10 s. Note that before the bad-segments definition, included segments lasting less than 5 seconds were also rejected. This decision was made because sandwiched periods (i.e., rejected-included-rejected) usually correspond to fully bad segments where the rejection algorithm did not mark all as bad). We call the included segments *tInc*. Afterward, we corrected motion artifacts by applying Spline interpolation (Scholkmann et al., 2010) using the Homer3 function hmrR_MotionCorrectSpline (p = 0.99), followed by Wavelet correction (Molavi & Dumont, 2012) using the Homer3 function hmrR_MotionCorrectWavelet (iqr = 1.5). Finally, we re-detected motion artifacts in the corrected data, and if new segments had more than 50 % of channels rejected, they were marked as bad in *tInc*. A final rejection matrix (*tIncCh*) was obtained based on the last artifacts detection, saturation, SNR<15, and SCI<0.6, and later used to reject specific channels from included blocks.

Subsequent steps of the analysis were performed in Python using MNE (version 1.7.0) and MNE-NIRS (version 0.6.0) (Luke et al., 2021). The data were band-pass filtered using an FIR filter between 0.01 and 0.3 Hz (transition bandwidths of 0.005 Hz for the high-pass and 0.1 Hz for low-pass) and converted to optical density using the modified Beer-Lambert law (partial path length factor 4.75; Scholkmann & Wolf, 2013). To obtain the HRF, data were segmented from -5 to 20 s relative to the onset of each stimulus block, linearly detrended, and baseline-corrected using the pre-stimulus interval. Channels for specific blocks were rejected if: (1) marked as bad during more than 50 % of the block in the rejection matrix *tIncCh*, (2) had an outlier peak-to-peak signal change defined as > *q*_3_ + 2 x (*q*_3 –_ *q*_1_), computed on normalized data across channels and blocks. Blocks were rejected if: (1) overlapped with not included segments (i.e., *tInc*=0), (2) had more than 35% of the active channels rejected. Subjects were rejected if more than 35% (more than 15 out of 42) of the channels were excluded from the recording (pruned channels). A testing sequence (familiarization/interference/test) for a given subject was excluded if fewer than five blocks were retained out of the 15 blocks (5 familiarization, 5 interference, 5 test). Of the 32 subjects with included data, 31 completed the same-word condition sequence, and 26 completed the novel-word condition sequence (25 both). On average, we obtained data for 22.3 subjects (range=[18, 27], std=2.56) for each experimental block. The average number of blocks included for each phase (out of 5) across subjects was, for the sequence with the same word: 4.1 (SD 1.1) target blocks, 3.9 (SD 1.2) interference blocks, and 3.5 (SD 1.4) test blocks; and for the sequence with the novel word: 4.2 (SD 1.1) target blocks, 3.8 (SD 1.3) interference blocks, and 4.0 (SD 1.1) test blocks. The mean percentage of channels rejected for the whole recording among the included subjects was 5.21% (SD 6.66; range [0, 30.95] %).

The channel data were combined into six symmetric ROIs: IFG (left and right, each comprising 4 channels), STG (left and right, each comprising 5 channels), and PL (left and right, each comprising 7 channels) (Figure 1A). On average across subjects and blocks, the activity for IFG left resulted from 4.0 channels (SD 0.16), IFG right 3.8 channels (SD 0.38), STG left 4.9 channels (SD 0.35), STG right 4.7 channels (SD 0.71), PL left 6.5 channels (SD 0.93), and PL right 5.4 channels (SD 1.58). The mean activity for each block over the time window [0, 15] s was used for statistical analysis. The time window was determined from the grand-average HRF across all blocks and subjects, which peaked at ∼7 s from stimulus onset and returned to baseline at ∼15 s (Supplementary Figure 1).

### Statistical analysis

Changes in the concentration of oxygenated hemoglobin (HbO) and deoxygenated hemoglobin (HbR) were calculated. We used LMM for the analysis, with the mean activation as the dependent variable. Fixed effects were nested within the block number and the ROIs, while the participant was included as a random effect. The models were solved in R (version 4.2.1) using the lme4 package (version 1.1.31).

## Acknowledgments

We would like to express our gratitude to the Neonatal Care Unit of the Unit of Neonatology and the Obstetric Division of the University Hospital of Padova for the recruitment of neonates and thank parents of newborns for their participation and cooperation.

## Funding information

This work was funded by the European Union (ERC-2021-STG, IN-MIND, Grant 101043216).

## Additional Information

### Data Availability Statement

The anonymized data collected are available as open data via the University of Padova online data repository: https://researchdata.cab.unipd.it/1403/ (DOI: 10.25430/researchdata.cab.unipd.it.00001403).

### Code Availability Statement

The code used for data preprocessing and analysis is available from the corresponding author upon request.

### Author Contributions

Conceptualisation: E.V., A.F., SB-V; Methodology: E.V., A.F., SB-V; Data collection: E.V., A.F., E.B.; Formal analysis: A.F.; Writing – original draft preparation; review & editing: E.V., A.F., E.B., SB-V; Supervision: SB-V; Project administration: SB-V; Funding acquisition: SB-V.

### Competing Interest Statement

The authors have declared that no competing interests exist.

## Notes

### Competing Interest Statement

The authors have declared no competing interest.

### Summary of Updates

Revisions have been made with particular attention to providing justification for the methods and extending the discussion.

## References

Agus, T. R., Paquette, S., Suied, C., Pressnitzer, D., & Belin, P. (2017). Voice selectivity in the temporal voice area despite matched low-level acoustic cues. Scientific Reports, 7(1), 11526. 10.1038/s41598-017-11684-1

Alberini, C. M., & Travaglia, A. (2017). Infantile Amnesia: A Critical Period of Learning to Learn and Remember. The Journal of Neuroscience, 37(24), 5783–5795. 10.1523/JNEUROSCI.0324-17.2017

Alexopoulos, J., Giordano, V., Doering, S., Seidl, R., Benavides-Varela, S., Russwurm, M., Greenwood, S., Berger, A., & Bartha-Doering, L. (2022). Sex differences in neural processing of speech in neonates. Cortex, 157, 117–128. 10.1016/j.cortex.2022.09.007

Alexopoulos, J., Giordano, V., Janda, C., Benavides-Varela, S., Seidl, R., Doering, S., Berger, A., & Bartha-Doering, L. (2021). The duration of intrauterine development influences discrimination of speech prosody in infants. Developmental Science, 24(5), e13110. 10.1111/desc.13110

Apfelbaum, K. S., & McMurray, B. (2011). Using Variability to Guide Dimensional Weighting: Associative Mechanisms in Early Word Learning. Cognitive Science, 35(6), 1105–1138. 10.1111/j.1551-6709.2011.01181.x

Bastianello, T., Nascimben, C., Loschiavo, A. G.-M. A., Reoyo-Serrano, N., Dassie, F., & Benavides-Varela, S. (2025). Infant sleep time: A comparative analysis of assessment methods. Sleep Medicine, 136, 106838. 10.1016/j.sleep.2025.106838

Behm, L., Turk-Browne, N. B., & Kibbe, M. M. (2025). The ubiquity of episodic-like memory during infancy. Trends in Cognitive Sciences, 29(11), 1034–1047. 10.1016/j.tics.2025.04.003

Belin, P., Zatorre, R. J., & Ahad, P. (2002). Human temporal-lobe response to vocal sounds. Cognitive Brain Research, 13(1), 17–26. 10.1016/S0926-6410(01)00084-2

Belin, P., Zatorre, R. J., Lafaille, P., Ahad, P., & Pike, B. (2000). Voice-selective areas in human auditory cortex. Nature, 403(6767), 309–312. 10.1038/35002078

Benavides-Varela, S. (2012). The Origin of Memory for Language. SISSA, Trieste – Italy

Benavides-Varela, S., & Gervain, J. (2017). Learning word order at birth: A NIRS study. Developmental Cognitive Neuroscience, 25, 198–208. 10.1016/j.dcn.2017.03.003

Benavides-Varela, S., Gómez, D. M., Macagno, F., Bion, R. A. H., Peretz, I., & Mehler, J. (2011). Memory in the Neonate Brain. PLoS ONE, 6(11), e27497. 10.1371/journal.pone.0027497

Benavides-Varela, S., Hochmann, J.-R., Macagno, F., Nespor, M., & Mehler, J. (2012). Newborn’s brain activity signals the origin of word memories. Proceedings of the National Academy of Sciences, 109(44), 17908–17913. 10.1073/pnas.1205413109

Blasi, A., Mercure, E., Lloyd-Fox, S., Thomson, A., Brammer, M., Sauter, D., Deeley, Q., Barker, G. J., Renvall, V., Deoni, S., Gasston, D., Williams, S. C. R., Johnson, M. H., Simmons, A., & Murphy, D. G. M. (2011). Early Specialization for Voice and Emotion Processing in the Infant Brain. Current Biology, 21(14), 1220–1224. 10.1016/j.cub.2011.06.009

Bodin, C., Takerkart, S., Belin, P., & Coulon, O. (2018). Anatomo-functional correspondence in the superior temporal sulcus. Brain Structure and Function, 223(1), 221–232. 10.1007/s00429-017-1483-2

Boersma, P., & van Heuven, V. (2001). Speak and unSpeak with PRAAT. Glot International, 5, 341–347.

Bouchon, C., Nazzi, T., & Gervain, J. (2015). Hemispheric Asymmetries in Repetition Enhancement and Suppression Effects in the Newborn Brain. PLOS ONE, 10(10), e0140160. 10.1371/journal.pone.0140160

Bunge, S. A. (2001). Prefrontal regions involved in keeping information in and out of mind. Brain, 124(10), 2074–2086. 10.1093/brain/124.10.2074

Cabeza, R., & Nyberg, L. (2000). Neural bases of learning and memory: Functional neuroimaging evidence: Current Opinion in Neurology, 13(4), 415–421. 10.1097/00019052-200008000-00008

Cheng, Y., Lee, S.-Y., Chen, H.-Y., Wang, P.-Y., & Decety, J. (2012). Voice and Emotion Processing in the Human Neonatal Brain. Journal of Cognitive Neuroscience, 24(6), 1411–1419. 10.1162/jocn_a_00214

Cheour-Luhtanen, M., Alho, K., Kujala, T., Sainio, K., Reinikainen, K., Renlund, M., Aaltonen, O., Eerola, O., & Näätänen, R. (1995). Mismatch negativity indicates vowel discrimination in newborns. Hearing Research, 82(1), 53–58. 10.1016/0378-5955(94)00164-L

Cooper, R. P., & Aslin, R. N. (1989). The language environment of the young infant: Implications for early perceptual development. Canadian Journal of Psychology / Revue Canadienne de Psychologie, 43(2), 247–265. 10.1037/h0084216

DeCasper, A. J., & Fifer, W. P. (1980). Of Human Bonding: Newborns Prefer Their Mothers’ Voices. Science, 208(4448), 1174–1176. 10.1126/science.7375928

Dehaene-Lambertz, G., & Baillet, S. (1998). A phonological representation in the infant brain: NeuroReport, 9(8), 1885–1888. 10.1097/00001756-199806010-00040

Dehaene-Lambertz, G., Dehaene, S., & Hertz-Pannier, L. (2002). Functional Neuroimaging of Speech Perception in Infants. Science, 298(5600), 2013–2015. 10.1126/science.1077066

Dehaene-Lambertz, G., & Pena, M. (2001). Electrophysiological evidence for automatic phonetic processing in neonates: Neuroreport, 12(14), 3155–3158. 10.1097/00001756-200110080-00034

DeWitt, I., & Rauschecker, J. P. (2012). Phoneme and word recognition in the auditory ventral stream. Proceedings of the National Academy of Sciences, 109(8). 10.1073/pnas.1113427109

Fecteau, S., Armony, J. L., Joanette, Y., & Belin, P. (2004). Priming of non-speech vocalizations in male adults: The influence of the speaker’s gender. Brain and Cognition, 55(2), 300–302. 10.1016/j.bandc.2004.02.024

Fernald, A., & Kuhl, P. (1987). Acoustic determinants of infant preference for motherese speech. Infant Behavior and Development, 10(3), 279–293. 10.1016/0163-6383(87)90017-8

Fló, A., Benjamin, L., Palu, M., & Dehaene-Lambertz, G. (2022). Sleeping neonates track transitional probabilities in speech but only retain the first syllable of words. Scientific Reports, 12(1), 4391. 10.1038/s41598-022-08411-w

Fló, A., Benjamin, L., Palu, M., & Dehaene-Lambertz, G. (2025). Statistical learning beyond words in human neonates. 10.7554/eLife.101802.2

Fló, A., Brusini, P., Macagno, F., Nespor, M., Mehler, J., & Ferry, A. L. (2019). Newborns are sensitive to multiple cues for word segmentation in continuous speech. Developmental Science, 22(4), e12802. 10.1111/desc.12802

Forgács, B., Tauzin, T., Gergely, G., & Gervain, J. (2022). The newborn brain is sensitive to the communicative function of language. Scientific Reports, 12(1), 1220. 10.1038/s41598-022-05122-0

Gervain, J., Macagno, F., Cogoi, S., Peña, M., & Mehler, J. (2008). The neonate brain detects speech structure. Proceedings of the National Academy of Sciences, 105(37), 14222–14227. 10.1073/pnas.0806530105

Giordano, V., Alexopoulos, J., Spagna, A., Benavides-Varela, S., Peganc, K., Kothgassner, O. D., Klebermass-Schrehof, K., Olischar, M., Berger, A., & Bartha-Doering, L. (2021). Accent discrimination abilities during the first days of life: An fNIRS study. Brain and Language, 223, 105039. 10.1016/j.bandl.2021.105039

Goldinger, S. D. (1996). Words and voices: Episodic traces in spoken word identification and recognition memory. *Journal of Experimental Psychology: Learning*, Memory, and Cognition, 22(5), 1166–1183. 10.1037/0278-7393.22.5.1166

Grossmann, T., Oberecker, R., Koch, S. P., & Friederici, A. D. (2010). The Developmental Origins of Voice Processing in the Human Brain. Neuron, 65(6), 852–858. 10.1016/j.neuron.2010.03.001

Halle, M. (1985). Speculations about the Representations of Words in Memory. In M. Halle, From Memory to Speech and Back (pp. 101–114). V. A. Fromkin.

Hickok, G., & Poeppel, D. (2007). The cortical organization of speech processing. Nature Reviews Neuroscience, 8(5), 393–402. 10.1038/nrn2113

Houston, D. M., & Jusczyk, P. W. (2000). The role of talker-specific information in word segmentation by infants. Journal of Experimental Psychology: Human Perception and Performance, 26(5), 1570–1582. 10.1037/0096-1523.26.5.1570

Huppert, T. J., Diamond, S. G., Franceschini, M. A., & Boas, D. A. (2009). HomER: A review of time-series analysis methods for near-infrared spectroscopy of the brain. Applied Optics, 48(10), D280. 10.1364/AO.48.00D280

Issard, C., & Gervain, J. (2018). Variability of the hemodynamic response in infants: Influence of experimental design and stimulus complexity. Developmental Cognitive Neuroscience, 33, 182–193. 10.1016/j.dcn.2018.01.009

Jusczyk, P. W. (1997). The discovery of spoken language (1st MIT Press pbk. ed). MIT Press.

Jusczyk, P. W., & Aslin, R. N. (1995). Infants′ Detection of the Sound Patterns of Words in Fluent Speech. Cognitive Psychology, 29(1), 1–23. 10.1006/cogp.1995.1010

Katus, L., Blasi, A., McCann, S., Mason, L., Mbye, E., Touray, E., Ceesay, M., De Haan, M., Moore, S. E., Elwell, C. E., & Lloyd-Fox, S. (2023). Longitudinal fNIRS and EEG metrics of habituation and novelty detection are correlated in 1–18-month-old infants. NeuroImage, 274, 120153. 10.1016/j.neuroimage.2023.120153

Kotilahti, K., Nissilä, I., Näsi, T., Lipiäinen, L., Noponen, T., Meriläinen, P., Huotilainen, M., & Fellman, V. (2010). Hemodynamic responses to speech and music in newborn infants. Human Brain Mapping, 31(4), 595–603. 10.1002/hbm.20890

Lee, C. W., Blanco, B., Dempsey, L., Chalia, M., Hebden, J. C., Caballero-Gaudes, C., Austin, T., & Cooper, R. J. (2020). Sleep State Modulates Resting-State Functional Connectivity in Neonates. Frontiers in Neuroscience, 14, 347. 10.3389/fnins.2020.00347

Lloyd-Fox, S., Blasi, A., McCann, S., Rozhko, M., Katus, L., Mason, L., Austin, T., Moore, S. E., Elwell, C. E., & The BRIGHT project team. (2019). Habituation and novelty detection fNIRS brain responses in 5- and 8-month-old infants: The Gambia and UK. Developmental Science, 22(5), e12817. 10.1111/desc.12817

Luke, R., Larson, E., Shader, M. J., Innes-Brown, H., Van Yper, L., Lee, A. K. C., Sowman, P. F., & McAlpine, D. (2021). Analysis methods for measuring passive auditory fNIRS responses generated by a block-design paradigm. Neurophotonics, 8(02). 10.1117/1.NPh.8.2.025008

Mahmoudzadeh, M., Dehaene-Lambertz, G., Fournier, M., Kongolo, G., Goudjil, S., Dubois, J., Grebe, R., & Wallois, F. (2013a). Syllabic discrimination in premature human infants prior to complete formation of cortical layers. Proceedings of the National Academy of Sciences, 110(12), 4846–4851. 10.1073/pnas.1212220110

Mahmoudzadeh, M., Dehaene-Lambertz, G., Fournier, M., Kongolo, G., Goudjil, S., Dubois, J., Grebe, R., & Wallois, F. (2013b). Syllabic discrimination in premature human infants prior to complete formation of cortical layers. Proceedings of the National Academy of Sciences, 110(12), 4846–4851. 10.1073/pnas.1212220110

Martinez-Alvarez, A., Benavides-Varela, S., Lapillonne, A., & Gervain, J. (2023). Newborns discriminate utterance-level prosodic contours. Developmental Science, 26(2), e13304. 10.1111/desc.13304

Mathias, S. R., & Von Kriegstein, K. (2014). How do we recognise who is speaking. *Frontiers in Bioscience*, S6(1), 92–109. 10.2741/S417

McClelland, J. L., & Elman, J. L. (1986). The TRACE model of speech perception. Cognitive Psychology, 18(1), 1–86. 10.1016/0010-0285(86)90015-0

Mehler, J., Bertoncini, J., Barriere, M., & Jassik-Gerschenfeld, D. (1978). Infant Recognition of Mother’s Voice. Perception, 7(5), 491–497. 10.1068/p070491

Mesgarani, N., Cheung, C., Johnson, K., & Chang, E. F. (2014). Phonetic Feature Encoding in Human Superior Temporal Gyrus. Science, 343(6174), 1006–1010. 10.1126/science.1245994

Molavi, B., & Dumont, G. A. (2012). Wavelet-based motion artifact removal for functional near-infrared spectroscopy. Physiological Measurement, 33(2), 259–270. 10.1088/0967-3334/33/2/259

Müller G. E. & Pilzecker A. (1900). Experimentelle Beiträge zur Lehre vom Gedächtnis. Zeitschrift Für Psychologie. Ergänzungsband, 1, 1–300.

Nakano, T., Watanabe, H., Homae, F., & Taga, G. (2009). Prefrontal Cortical Involvement in Young Infants’ Analysis of Novelty. Cerebral Cortex, 19(2), 455–463. 10.1093/cercor/bhn096

Norris, D., McQueen, J. M., & Cutler, A. (2000). Merging information in speech recognition: Feedback is never necessary. Behavioral and Brain Sciences, 23(3), 299–325. 10.1017/S0140525X00003241

Novick, J. M., Trueswell, J. C., & Thompson-Schill, S. L. (2010). Broca’s Area and Language Processing: Evidence for the Cognitive Control Connection. Language and Linguistics Compass, 4(10), 906–924. 10.1111/j.1749-818X.2010.00244.x

Nygaard, L. C., Sommers, M. S., & Pisoni, D. B. (1994). Speech Perception as a Talker-Contingent Process. Psychological Science, 5(1), 42–46. 10.1111/j.1467-9280.1994.tb00612.x

Palmeri, T. J., Goldinger, S. D., & Pisoni, D. B. (1993). Episodic encoding of voice attributes and recognition memory for spoken words. *Journal of Experimental Psychology: Learning*, Memory, and Cognition, 19(2), 309–328. 10.1037/0278-7393.19.2.309

Peirce, J., Gray, J. R., Simpson, S., MacAskill, M., Höchenberger, R., Sogo, H., Kastman, E., & Lindeløv, J. K. (2019). PsychoPy2: Experiments in behavior made easy. Behavior Research Methods, 51(1), 195–203. 10.3758/s13428-018-01193-y

Peña, M., Maki, A., KovaciJić, D., Dehaene-Lambertz, G., Koizumi, H., Bouquet, F., & Mehler, J. (2003). Sounds and silence: An optical topography study of language recognition at birth. Proceedings of the National Academy of Sciences, 100(20), 11702–11705. 10.1073/pnas.1934290100

Perani, D., Saccuman, M. C., Scifo, P., Spada, D., Andreolli, G., Rovelli, R., Baldoli, C., & Koelsch, S. (2010). Functional specializations for music processing in the human newborn brain. Proceedings of the National Academy of Sciences, 107(10), 4758–4763. 10.1073/pnas.0909074107

Peretz, I., Vuvan, D., Lagrois, M.-É., & Armony, J. L. (2015). Neural overlap in processing music and speech. Philosophical Transactions of the Royal Society B: Biological Sciences, 370(1664), 20140090. 10.1098/rstb.2014.0090

Pernet, C. R., McAleer, P., Latinus, M., Gorgolewski, K. J., Charest, I., Bestelmeyer, P. E. G., Watson, R. H., Fleming, D., Crabbe, F., Valdes-Sosa, M., & Belin, P. (2015). The human voice areas: Spatial organization and inter-individual variability in temporal and extra-temporal cortices. NeuroImage, 119, 164–174. 10.1016/j.neuroimage.2015.06.050

Pisoni, D. B., & Luce, P. A. (1987). Acoustic-phonetic representations in word recognition. Cognition, 25(1–2), 21–52. 10.1016/0010-0277(87)90003-5

Pollonini, L., Olds, C., Abaya, H., Bortfeld, H., Beauchamp, M. S., & Oghalai, J. S. (2014). Auditory cortex activation to natural speech and simulated cochlear implant speech measured with functional near-infrared spectroscopy. Hearing Research, 309, 84–93. 10.1016/j.heares.2013.11.007

Rankin, C. H., Abrams, T., Barry, R. J., Bhatnagar, S., Clayton, D. F., Colombo, J., Coppola, G., Geyer, M. A., Glanzman, D. L., Marsland, S., McSweeney, F. K., Wilson, D. A., Wu, C.-F., & Thompson, R. F. (2009). Habituation revisited: An updated and revised description of the behavioral characteristics of habituation. Neurobiology of Learning and Memory, 92(2), 135–138. 10.1016/j.nlm.2008.09.012

Scholkmann, F., Spichtig, S., Muehlemann, T., & Wolf, M. (2010). How to detect and reduce movement artifacts in near-infrared imaging using moving standard deviation and spline interpolation. Physiological Measurement, 31(5), 649–662. 10.1088/0967-3334/31/5/004

Scholkmann, F., & Wolf, M. (2013). General equation for the differential pathlength factor of the frontal human head depending on wavelength and age. Journal of Biomedical Optics, 18(10), 105004. 10.1117/1.JBO.18.10.105004

Schönwiesner, M., Rübsamen, R., & Von Cramon, D. Y. (2005). Spectral and Temporal Processing in the Human Auditory Cortex—Revisited. Annals of the New York Academy of Sciences, 1060(1), 89–92. 10.1196/annals.1360.051

Simon, S., Lazeyras, F., Sigrist, A.-D., Ecoffey, M., Guatieri, S., Van De Ville, D., Borradori-Tolsa, C., Pelizzone, M., & Hüppi, P. (2009). FMRI study of newborn perception of mother’s voice: A comparative study of premature infants at term age and term born neonates. NeuroImage, 47, S54. 10.1016/S1053-8119(09)70181-8

Singh, L. (2008). Influences of high and low variability on infant word recognition. Cognition, 106(2), 833–870. 10.1016/j.cognition.2007.05.002

Singh, L., Morgan, J. L., & White, K. S. (2004). Preference and processing: The role of speech affect in early spoken word recognition. Journal of Memory and Language, 51(2), 173–189. 10.1016/j.jml.2004.04.004

Spence, M. J., & Freeman, M. S. (1996). Newborn infants prefer the maternal low-pass filtered voice, but not the maternal whispered voice. Infant Behavior and Development, 19(2), 199–212. 10.1016/S0163-6383(96)90019-3

Telkemeyer, S., Rossi, S., Koch, S. P., Nierhaus, T., Steinbrink, J., Poeppel, D., Obrig, H., & Wartenburger, I. (2009). Sensitivity of Newborn Auditory Cortex to the Temporal Structure of Sounds. The Journal of Neuroscience, 29(47), 14726–14733. 10.1523/JNEUROSCI.1246-09.2009

Thompson-Schill, S. L., D’Esposito, M., Aguirre, G. K., & Farah, M. J. (1997). Role of left inferior prefrontal cortex in retrieval of semantic knowledge: A reevaluation. Proceedings of the National Academy of Sciences, 94(26), 14792–14797. 10.1073/pnas.94.26.14792

Trainor, L. J., Clark, E. D., Huntley, A., & Adams, B. A. (1997). The acoustic basis of preferences for infant-directed singing. Infant Behavior and Development, 20(3), 383–396. 10.1016/S0163-6383(97)90009-6

Tulving, E. (1993). What Is Episodic Memory? Current Directions in Psychological Science, 2(3), 67–70. 10.1111/1467-8721.ep10770899

Uchitel, J., Blanco, B., Collins-Jones, L., Edwards, A., Porter, E., Pammenter, K., Hebden, J., Cooper, R. J., & Austin, T. (2023). Cot-side imaging of functional connectivity in the developing brain during sleep using wearable high-density diffuse optical tomography. NeuroImage, 265, 119784. 10.1016/j.neuroimage.2022.119784

Van Heugten, M., Bergmann, C., & Cristia, A. (2015). The effects of talker voice and accent on young children’s speech perception. In Individual Differences in Speech Production and Perception (pp. 57–88). In S. Fuchs, D. Pape, C. Petrone, & P. Perrier (Eds.).

Werker, J. F., & Curtin, S. (2005). PRIMIR: A Developmental Framework of Infant Speech Processing. Language Learning and Development, 1(2), 197–234. 10.1080/15475441.2005.9684216

Wildgruber, D., Ackermann, H., Kreifelts, B., & Ethofer, T. (2006). Cerebral processing of linguistic and emotional prosody: fMRI studies. In Progress in Brain Research (Vol. 156, pp. 249–268). Elsevier. 10.1016/S0079-6123(06)56013-3

Yates, T. S., Fel, J., Choi, D., Trach, J. E., Behm, L., Ellis, C. T., & Turk-Browne, N. B. (2025). Hippocampal encoding of memories in human infants. Science, 387(6740), 1316–1320. 10.1126/science.adt7570

Zatorre, R. J., & Belin, P. (2001). Spectral and Temporal Processing in Human Auditory Cortex. Cerebral Cortex, 11(10), 946–953. 10.1093/cercor/11.10.946

Zatorre, R. J., Belin, P., & Penhune, V. B. (2002). Structure and function of auditory cortex: Music and speech. Trends in Cognitive Sciences, 6(1), 37–46. 10.1016/S1364-6613(00)01816-7

