## Supplementary material for "Verbal Episodic Processing in Newborns": SI_eLife.pdf

### Supplementary materials

#### Methods

**Supplementary Table 1: Acoustic features across speakers.** Analysis of acoustic features across speakers (familiarization and test pseudowords: voice_female_1; voice_female_2; interference words: voice_male_1; voice_male_2). Intensity and duration were held constant for each word (70dB and 700ms respectively) while pitch (F0) and timbre (F1 and F2 of vowels) were extracted to assess acoustic differences across speakers.

| **mita** | |
|---|---|
| *voice_female_1* | *voice_female_1* |
| pitch: 201.57<br>F1_mita_/i/: 376.65<br>F2_mita_/i/: 1249.95<br>F1_mita_/a/: 875.63<br>F2_mita_/a/: 1499.28<br>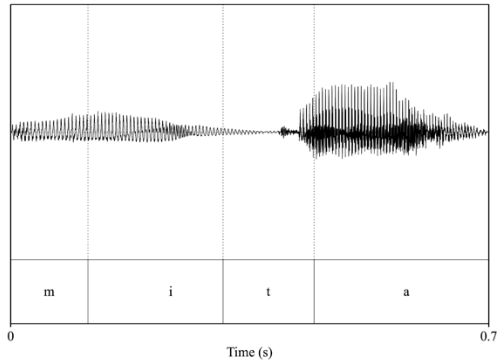<br> | pitch: 188.43<br>F1_mita_/i/: 360.56<br>F2_mita_/i/: 2732.94<br>F1_mita_/a/: 865.07<br>F2_mita_/a/: 1369.88<br>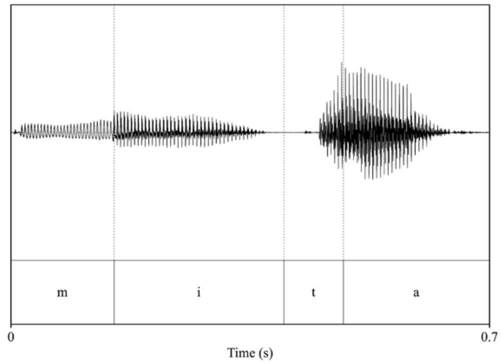<br> |
| **pelu** | |
| *voice_female_1* | *voice_female_2* |
| pitch: 171.44<br>F1_pelu_/e/: 413.5<br>F2_pelu_/e/: 2441.28<br>F1_pelu_/u/: 405.49<br>F2_pelu_/u/: 822.86 | pitch: 179.17<br>F1_pelu_/e/: 385.75<br>F2_pelu_/e/: 2480.84<br>F1_pelu_/u/: 376.49<br>F2_pelu_/u/: 837.6 |

<table>
<tr><td>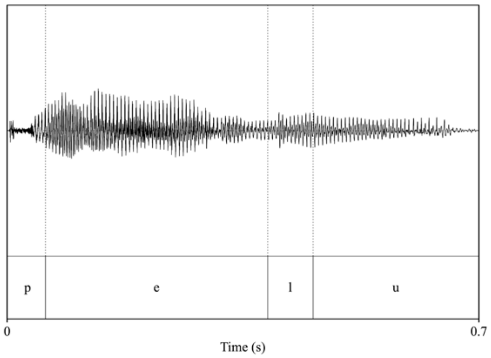
</td><td>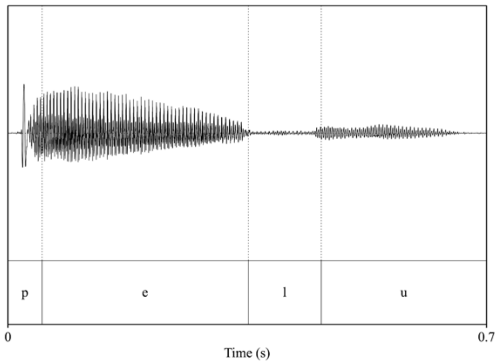
</td></tr>
<tr><td colspan="2">voli</td></tr>
<tr><td>voice_female_1</td><td>voice_female_2</td></tr>
<tr><td>pitch: 193.75<br>F1_voli_/o/: 457.55<br>F2_voli_/o/: 908.18<br>F1_voli_/i/: 372.43<br>F2_voli_/i/: 2521.58<br>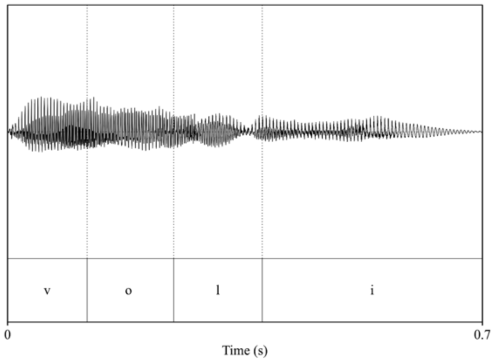
</td><td>pitch: 173.96<br>F1_voli_/o/: 433.28<br>F2_voli_/o/: 870.12<br>F1_voli_/i/: 349.75<br>F2_voli_/i/: 2691.6<br>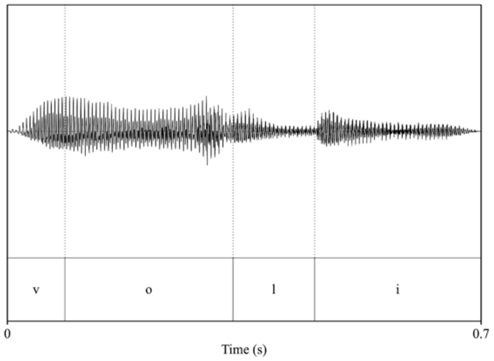
</td></tr>
<tr><td colspan="2">dafo</td></tr>
<tr><td>voice_male_1</td><td>voice_male_2</td></tr>
<tr><td>pitch: 84.89<br>F1_dafo_/a/: 807.9<br>F2_dafo_/a/: 1126.4<br>F1_dafo_/o/: 551.35<br>F2_dafo_/o/: 1339.42</td><td>pitch: 86.12<br>F1_dafo_/a/: 676.91<br>F2_dafo_/a/: 1268.19<br>F1_dafo_/o/: 502.94<br>F2_dafo_/o/: 774.73</td></tr>
</table>

| | |
|---|---|
| 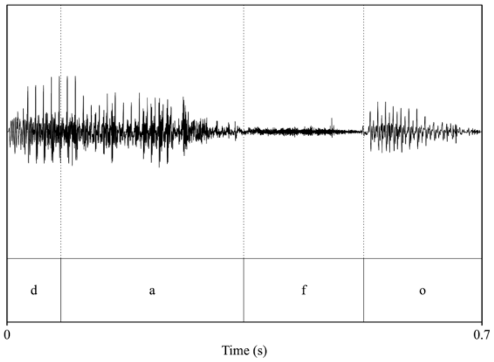 | 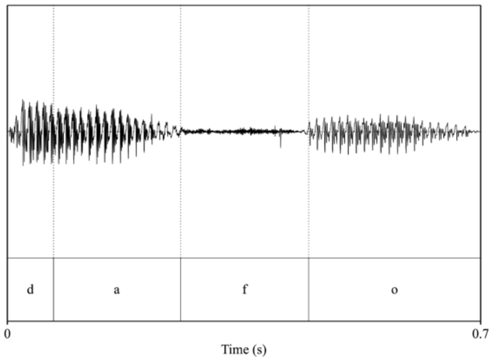 |
| **noke** | |
| *voice_male_1* | *voice_male_2* |
| pitch: 85.92<br>F1_noke_/o/: 604<br>F2_noke_/o/: 1017.2<br>F1_noke_/e/: 488.63<br>F2_noke_/e/: 1901.05<br>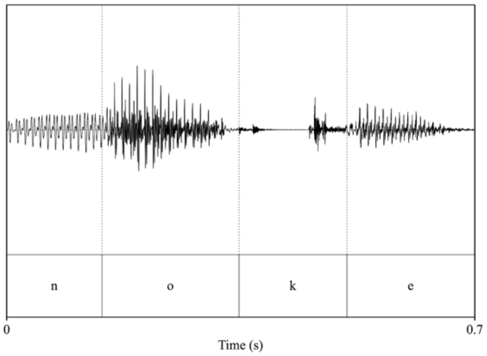 | pitch: 85.5<br>F1_noke_/o/: 593.86<br>F2_noke_/o/: 1018.64<br>F1_noke_/e/: 427.31<br>F2_noke_/e/: 1992.47<br>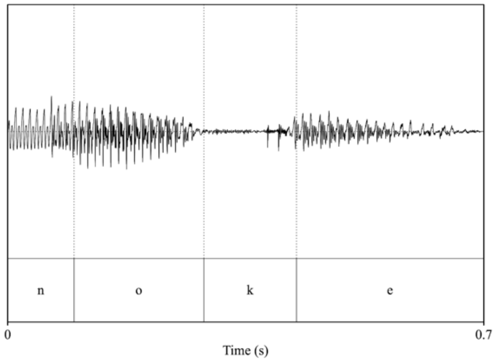 |

#### Results

##### HRF response across blocks.

We obtained the average response across all blocks in each ROI. This average response was used to define the time window for computing the average activity for each block. Since the grand average HRF returned to baseline level at ~15 s, we define the time window as [0, 15] s (Supplementary Figure 1).

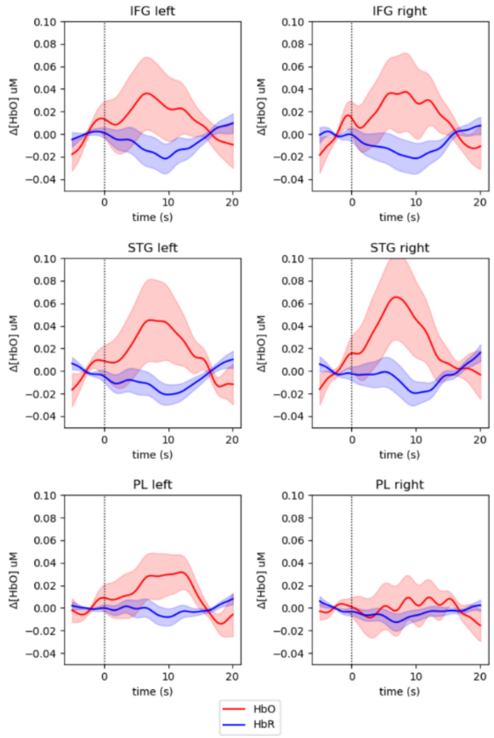


**Supplementary Figure 1: Grand HRF across all blocks**. Shaded areas represent the standard error across subjects. HbO is represented in red and HbR in blue.

#### Results for HbR.

**Habituation and novelty effects.** The LMM act ~-1+block:ROI + (1 | sub) during the learning phase, testing for activation differing from zero. There was deactivation in block 2 within left IFG ($\beta$=-0.0894, SE=0.025, $p$ = 0.0004). During the interference phase, it showed deactivation in block 2 over right IFG ($\beta$=-0.0987, SE=0.030, $p$ = 0.001) and activation in block 5 over left IFG ($\beta$=0.0686, SE=0.029, $p$ = 0.017).

The LMM act ~-1+ROI +ROI:blocknumber+ (1 | sub), with blocknumber ranging from 0 to 4, showed no significant linear changes during the learning phase ($p$>0.05). Instead during the interference phase the activity increased over left IFG (intercept=-0.0663, SE=0.030, p=0.027; slope=0.0217, SE=0.0082, p=0.0083), right IFG (intercept=-0.0946, SE=0.030, p=0.0019; slope=0.0253, SE=0.0082, p=0.0021), left STG (intercept=-0.0602, SE=0.030, p=0.047; slope=0.0197, SE=0.0082, p=0.017), and right PL (intercept=-0.0240, SE=0.031, p=0.4; slope=0.0192, SE=0.0083, p=0.020).

Finally, no differences were observed between the activity of the last block of familiarization and the first block of interference ($p$>0.05).

**Word recognition.** The LMM act ~-1+block:ROI +block:ROI:condition+ (1 | sub) showed no significant differences between conditions for any block and ROI ($p$>0.1).

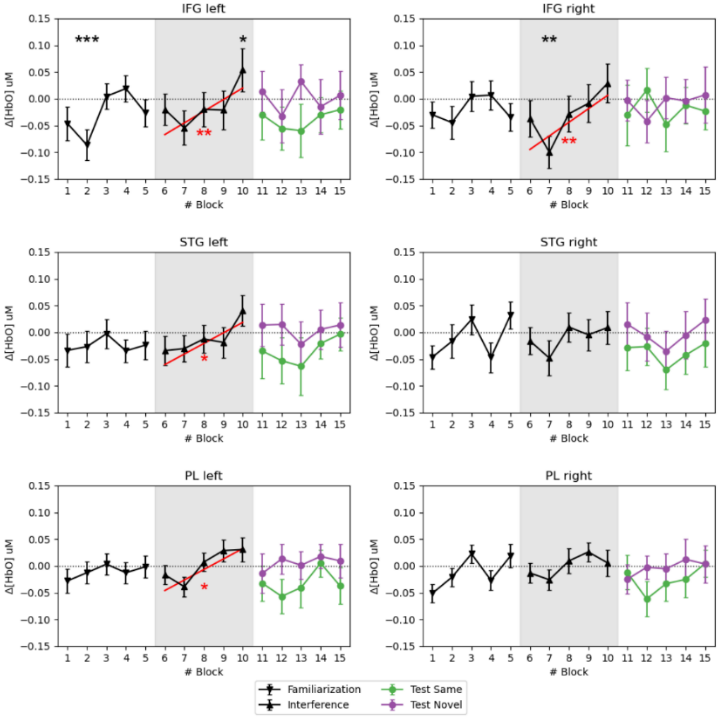

**Supplementary Figure 2: Block activity for HbR during the learning, interference, and test phases.** Error bars represent the standard errors. The black continuous line depicts responses averaged across all participants and conditions. The same-word condition (green) and the novel-word condition (purple) are plotted in the test phase. The black asterisks during the learning and interference phases indicate that the response differed from zero. The red lines indicate a significant linear trend, as indicated by the red asterisks. No significant differences between conditions were observed ($p>0.1$).

#### Description of the LMM used for testing for order effects

Activation patterns might differ from the first to the second sequence because infants retain information from the first sequence that affects the processing of the second or due to changes in cognitive state. Groups A heard first the sequence [X y X] (*same-word* condition) and then [X y Z] (*novel-word* condition), while groups B heard [X y Z] followed by [X y X]. Thus, sequences and conditions are inversely linked in each group.

During the learning and interference phases, we were interested in investigating whether (1) there were main differences due to the position in time of the phases (first sequence presentation or second), (2) there were differences between the groups in sequence 1, and (3) there were differences between the groups in sequence 2. Differences between groups within each sequence might be due to individual differences. In the case of the second sequence, they might also be related to the different content that the two groups heard in the first sequence. To do so, we run an LMM with sequence number and group as fixed effects and custom contrasts to produce interpretable results and test these hypotheses. The contrasts used were:

*(H0.1) main effect of sequence:*

0 = (groupA_sequence1 + groupA_sequence1 - groupB_sequence2 - groupB_sequence2)/2,

*(H0.2) differences between groups within the first sequence:*

0 = groupA_sequence1 - groupB_sequence1, and

*(H0.3) differences between groups within the second sequence:*

0 = groupA_sequence2 - groupB_sequence2.

The contrasts were nested within blocks and ROIs. The model was: act ~-1+block:ROI +block:ROI:C+ (1 | sub), where C corresponds to the contrasts. Therefore, a significant coefficient for contrast 1 signifies a main effect of sequence, while a significant coefficient for contrasts 2 and 3 implies differences between groups in sequences 1 and 2, respectively.

In the test phase, the group determines the conditions the participants hear in each sequence. Thus, we included the main effect of the condition in the LMM as contrasts of interest and added the differences between groups within each condition. Differences within conditions might, therefore, result from individual differences (group) or the test word appearing in the first or second sequence. The contrasts used were:

*(H0.1) main effect of condition:*

0 = (groupA_novel + groupB_novel - groupA_same - groupB_same)/2,

*(H0.2) differences between groups within the same-word condition:*

0 = groupA_same - groupB_same, and

*(H0.3) differences between groups within the novel-word condition:*

0 = groupA_novel - groupB_novel.

The contrasts were nested within blocks and ROIs. The model was: act ~-1+block:ROI +block:ROI:C+ (1 | sub), where C corresponds to the contrasts.

#### Activation patterns during learning.

Supplementary Figure 3 shows the activation patterns during the learning phase. Results are described in the main text.

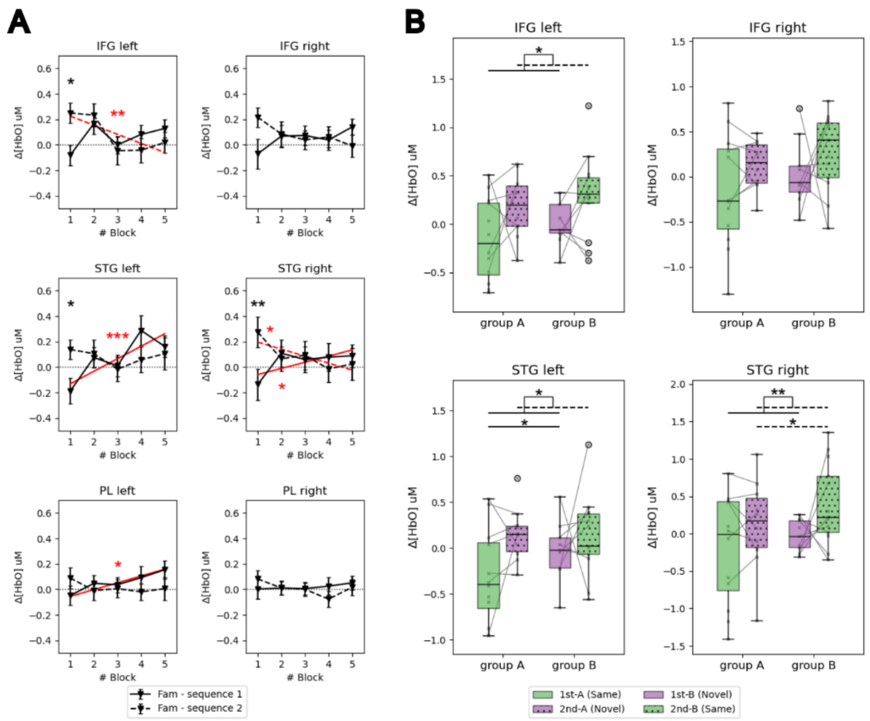


**Supplementary Figure 3: Differences in habituation responses during the familiarization phase across sequences. (A)** Mean activity for HbO over blocks during the first (full-line) and second (dashed-line) familiarization phases. Error bars represent the standard errors. Asterisks indicate blocks showing significant differences between the first and second familiarization phases. The red lines indicate a significant linear trend, as indicated by the red asterisks. For the first familiarization the LMM *act ~-1+ROI +ROI:blocknumber +(1|sub)*, with *blocknumber* ranging from 0 to 4 showed a significant increase in activity over blocks in the left and right STG (left: slope=0.098, SE=0.024, p=0.00005; right: slope=0.048, SE=0.024, p=0.045) and left PL (slope=0.053, SE=0.024, p=0.027). During the second familiarization, the activity was higher than zero in the first block and decreased with block number on the right STG (intercept=0.196, SE=0.071, p = 0.0066; slope=-0.055, SE=0.026, p=0.033), and left and right IFG (left: intercept=0.225, SE=0.071, p = 0.0019; slope=-0.071, SE=0.026, p=0.0056). **(B)** Mean activity for HbO during the first block of the familiarization phase over STG and IFG, where major differences between the first and second testing sequence were observed. First sequence: full box; second sequence: dotted box. The colors indicate whether the familiarization belongs to the same or novel word testing sequence.

#### Activation patterns during interference.

The LMM *act ~-1+block:ROI+(1|sub)* during the interference phase showed no significant differences ($p>0.05$), meaning activity did not significantly differ from zero in any block. The LMM testing for linear changes in activity *act ~-1+ROI+ROI:blocknumber+(1|sub)*, with *blocknumber*, showed a positive intercept with a negative slope in the right IFG (intercept=0.153, SE=0.0684, p=0.026; slope=-0.0413, SE=0.019, p=0.027), denoting initial high activity that decreased across blocks. Lastly, to investigate potential novelty effects arising from the transition from the learning to the interference phase, we compare the activity of the last block of the learning phase and the first block of interference phase. No significant differences were observed in any ROI ($p>0.1$).

The analysis investigating order effects during the interference phase showed higher activity during the second sequence in the third block over the left IFG ($\beta=-0.314$, SE=0.131, $p = 0.016$). No significant differences were observed between groups in the first or second sequences ($p>0.05$).

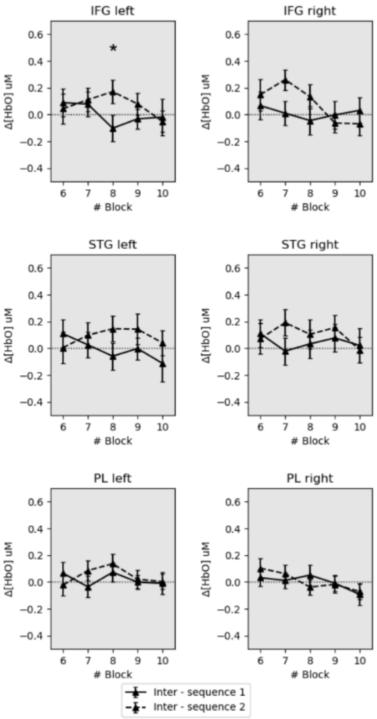


**Supplementary Figure 4: Habituation responses during the interference phase across sequences.** Mean activity for HbO over blocks during the first (full-line) and second (dashed-line) interference phases. Error bars represent the standard errors. Asterisks indicate blocks showing significant differences between the first and second familiarization phases.

#### Analysis at the channel level.

To ensure that the selection of the ROIs did not have a substantial influence on the results, we ran the analysis testing for differences between conditions during the test phase at the channel level. To do so, we run the model act~-1+block:channel+block:channel:condition+(1|sub). The significant coefficients contrasting the conditions are shown in Supplementary Table 2.

**Supplementary Table 2: Results for the analysis at the channel level, evaluating differences between conditions during the Test phase.** LMM: act~-1+block:channel+block:channel:condition+(1|sub). Only significant coefficients ($p<0.05$) for the contrast between conditions are shown.

| | Estimate | Std. Error | df | t value | Pr(>\|t\|) |
|---|---|---|---|---|---|
| block2:cha_S12_D12 | 0.409779 | 0.177042 | 7838.167 | 2.314586 | 0.020661 |
| block5:cha_S12_D12 | -0.55653 | 0.167365 | 7838.71 | -3.32523 | 0.000888 |
| block1:cha_S12_D13 | -0.50074 | 0.165594 | 7837.931 | -3.02393 | 0.002503 |
| block5:cha_S12_D13 | -0.36341 | 0.163359 | 7838.626 | -2.22462 | 0.026135 |
| block2:cha_S14_D12 | 0.500881 | 0.180004 | 7838.163 | 2.782615 | 0.005405 |
| block5:cha_S14_D12 | -0.43254 | 0.174754 | 7838.85 | -2.47513 | 0.01334 |
| block5:cha_S14_D13 | -0.32359 | 0.164814 | 7838.501 | -1.96335 | 0.049641 |
| block2:cha_S14_D14 | 0.470902 | 0.184399 | 7838.162 | 2.553705 | 0.010677 |
| block2:cha_S2_D1 | 0.431107 | 0.176339 | 7838.231 | 2.444757 | 0.014517 |
| block2:cha_S2_D2 | 0.354384 | 0.170551 | 7837.98 | 2.077884 | 0.037753 |
| block2:cha_S2_D3 | 0.382112 | 0.180017 | 7838.239 | 2.122645 | 0.033815 |
| block1:cha_S2_D4 | -0.34561 | 0.163692 | 7837.839 | -2.11136 | 0.034773 |
| block2:cha_S2_D4 | 0.50764 | 0.170551 | 7837.98 | 2.976476 | 0.002925 |
| block2:cha_S4_D4 | 0.460346 | 0.170551 | 7837.98 | 2.699176 | 0.006966 |
| block2:cha_S6_D4 | 0.536299 | 0.175077 | 7837.872 | 3.063223 | 0.002197 |
| block2:cha_S6_D5 | 0.590803 | 0.172379 | 7838.074 | 3.427358 | 0.000613 |
| block2:cha_S7_D5 | 0.420251 | 0.170551 | 7837.98 | 2.464085 | 0.013758 |
| block2:cha_S8_D6 | 0.404044 | 0.170551 | 7837.98 | 2.369059 | 0.017858 |
| block2:cha_S8_D7 | 0.360462 | 0.170551 | 7837.98 | 2.113519 | 0.034588 |
